# A *k*-mer-based method for the identification of phenotype-associated genomic biomarkers and predicting phenotypes of sequenced bacteria

**DOI:** 10.1101/302026

**Authors:** Erki Aun, Age Brauer, Veljo Kisand, Tanel Tenson, Maido Remm

**Affiliations:** Department of Bioinformatics, Institute of Molecular and Cell Biology, University of Tartu, Estonia; Institute of Technology, University of Tartu, Estonia

## Abstract

We have developed an easy-to-use and memory-efficient method called PhenotypeSeeker that (a) generates a *k*-mer-based statistical model for predicting a given phenotype and (b) predicts the phenotype from the sequencing data of a given bacterial isolate. The method was validated on 167 *Klebsiella pneumoniae* isolates (virulence), 200 *Pseudomonas aeruginosa* isolates (ciprofloxacin resistance) and 460 *Clostridium difficile* isolates (azithromycin resistance). The phenotype prediction models trained from these datasets performed with 88% accuracy on the *K. pneumoniae* test set, 88% on the *P. aeruginosa* test set and 96.5% on the *C. difficile* test set. Prediction accuracy was the same for assembled sequences and raw sequencing data; however, building the model from assembled genomes is significantly faster. On these datasets, the model building on a mid-range Linux server takes approximately 3 to 5 hours per phenotype if assembled genomes are used and 10 hours per phenotype if raw sequencing data are used. The phenotype prediction from assembled genomes takes less than one second per isolate. Thus, PhenotypeSeeker should be well-suited for predicting phenotypes from large sequencing datasets.

PhenotypeSeeker is implemented in Python programming language, is open-source software and is available at GitHub (https://github.com/bioinfo-ut/PhenotypeSeeker/).

**Summary:** Predicting phenotypic properties of bacterial isolates from their genomic sequences has numerous potential applications. A good example would be prediction of antimicrobial resistance and virulence phenotypes for use in medical diagnostics. We have developed a method that is able to predict phenotypes of interest from the genomic sequence of the isolate within seconds. The method uses statistical model that can be trained automatically on isolates with known phenotype. The method is implemented in Python programming language and can be run on low-end Linux server and/or on laptop computers.

## Introduction

The falling cost of sequencing has made genome sequencing affordable to a large number of labs, and therefore, there has been a dramatic increase in the number of genome sequences available for comparison in the public domain [1]. These developments have facilitated the genomic analysis of bacterial isolates. An increasing amount of bacterial whole genome sequencing (WGS) data has led to more and more genome-wide studies of DNA variation related to different phenotypes [2–7]. Among these studies, antibiotic resistance phenotypes are the most concerning and have garnered high public interest, especially since several multidrug-resistant strains have emerged worldwide [8]. The detection of known resistance-causing mutations as well as the search for new candidate biomarkers leading to resistance phenotypes requires reasonably rapid and easily applicable tools for processing and comparing the sequencing data of hundreds of isolated strains. However, there is still a lack of user-friendly software tools for the identification of genomic biomarkers from large sequencing datasets of bacterial isolates [9,10].

Methods that are based on sequence alignment are limited because they are strongly dependent on the availability of the list of previously described and confirmed resistance genes and mutations. New variations relevant to a bacterial phenotype would be missed if we rely on known markers. In addition, many bacterial species have extensive intra-species variation from small sequence-based differences to the absence or presence of whole genes or gene clusters. Choosing only one genome as a reference for searching for the variable components would be highly limiting.

*K*-mers, which are short DNA oligomers with length *k*, enable us to simultaneously discover a large set of single nucleotide variations, insertions and deletions associated with the phenotypes under study. The advantage of using *k*-mer-based methods in genomic biomarker discovery is that they do not require sequence alignments and can even be applied to raw sequencing data. Several *k*-mer-based tools for detecting the biomarkers behind different bacterial phenotypes have been previously published. The SEER program takes either a discrete or continuous phenotype as an input, counts variable-length *k*-mers and corrects for the clonal population structure [11]. SEER is a complex pipeline requiring several separate steps for the user to execute and currently has many system-level dependencies for successful compilation and installation. Another similar tool, Kover, handles only discrete phenotypes, counts user-defined size *k*-mers and does not use any correction for population structure [12]. The Neptune software targets so-called ‘signatures’ differentiating two groups of sequences but cannot locate smaller mutations, such as single isolated nucleotide variations. The ‘signatures’ that Neptune detects are relatively large genomic loci, which may include genomic islands, phage regions or operons [13].

We created PhenotypeSeeker as we observed the need for a tool that could combine all the benefits of the programs available but at the same time would be easily executable and would take a reasonable amount of computing resources without the need for dedicated high-performance computer hardware.

## Results

### Implementation

PhenotypeSeeker consist of two subprograms: ‘PhenotypeSeeker modeling’ and ‘PhenotypeSeeker prediction’. ‘PhenotypeSeeker modeling’ takes either assembled contigs or raw-read data as an input and builds a statistical model for phenotype prediction. The method starts with counting all possible *k*-mers from the input genomes, using the GenomeTester4 software package [14], followed by *k*-mer filtering by their frequency in strains. Subsequently, the *k*-mer selection for regression analysis is performed. In this step, to test the *k*-mers’ association with the phenotype, the method applies Welch’s two-sample t-test if the phenotype is continuous and a chi-squared test if it is binary. Finally, the logistic regression or linear regression model is built. The PhenotypeSeeker output gives the regression model in a binary format and three text files, which include the following: (1) the results of association tests, (2) the coefficients of *k*-mers in the regression model, (3) a FASTA file with phenotype-specific *k*-mers, assembled to longer contigs when possible, and (4) a summary of the regression analysis performed (Fig 1). Optionally, it is possible to use weighting for the strains to take into account the clonal population structure. The weights are based on a distance matrix of strains made with an alignment-free *k*-mer-based method called Mash [15]. The weights of each genome are calculated using the Gerstein, Sonnhammer and Cothia method [16]. ‘PhenotypeSeeker prediction’ uses the regression model generated by ‘PhenotypeSeeker modeling’ to conduct fast phenotype predictions on input samples (Fig 1). Using gmer_counter from the FastGT package [17], the tool searches the samples only for the *k*-mers used as parameters in the regression model. Predictions are then made based on the presence or absence of these *k*-mers.

**Fig 1.**
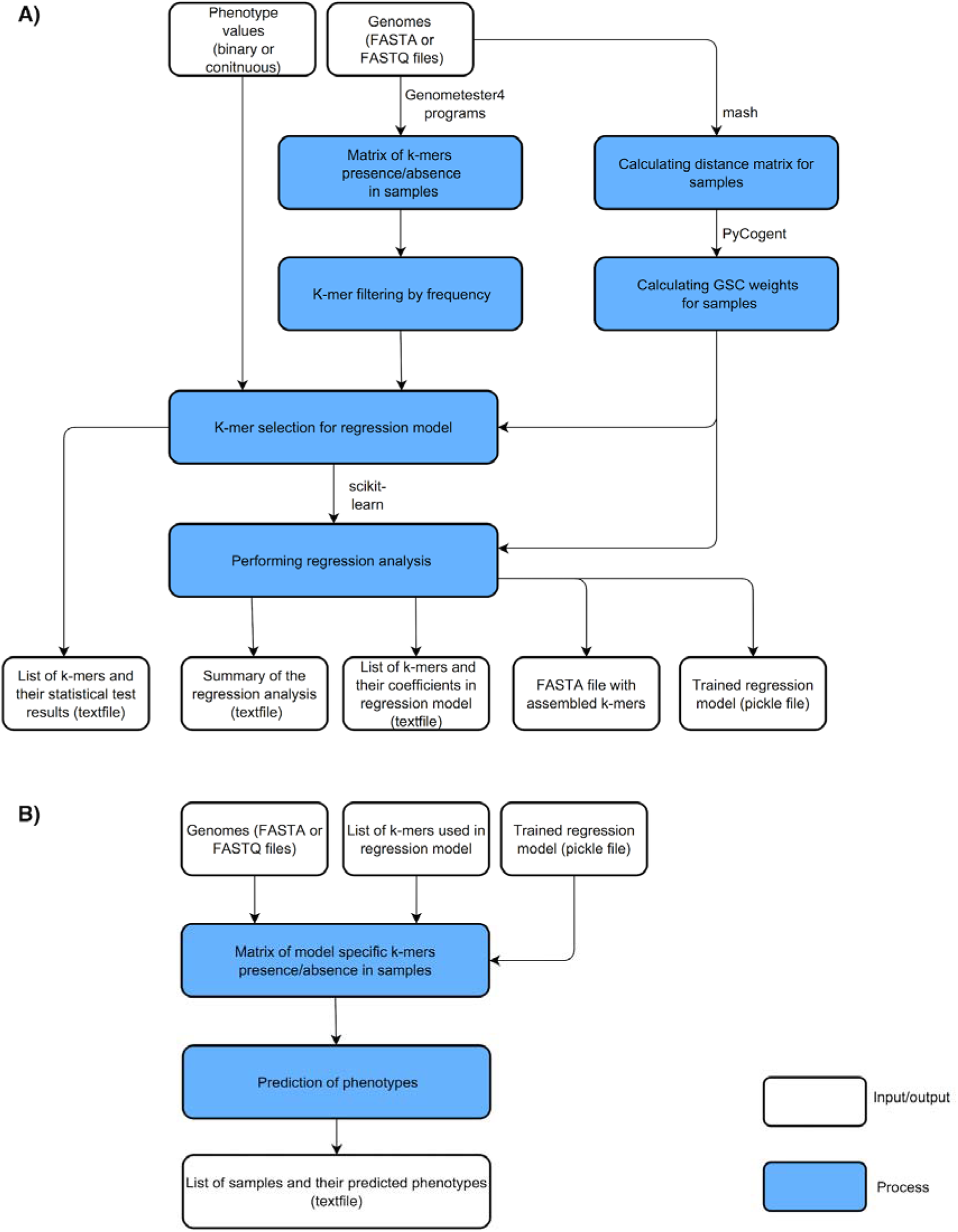
Schematic presentation of PhenotypeSeeker workflow. Panel A shows the ‘PhenotypeSeeker modeling’ steps, which generate the phenotype prediction model based on the input genomes and their phenotype values. Panel B shows the ‘PhenotypeSeeker prediction’ steps, which use the previously generated model to predict the phenotypes for input genomes.

PhenotypeSeeker uses fixed-length k-mers in all analyses. Thus, the *k*-mer length is an important factor influencing the overall software performance. The effects of *k*-mer length on speed, memory usage and accuracy were tested on a *P. aeruginosa* ciprofloxacin dataset. A general observation from that analysis is that the CPU time and the PhenotypeSeeker memory usage increase when the *k*-mer length increases (Fig 2). Previously described mutations in the *P. aeruginosaparC* and *gyrA* genes were always detected if the *k*-mer length was at least 13 nucleotides. We assume that in most cases, a *k*-mer length of 13 is sufficient to detect biologically relevant mutations, although in certain cases, longer *k*-mers might provide additional sensitivity. The k-mer length in PhenotypeSeeker is a user-selectable parameter. Although most of the phenotype detection can be performed with the default *k*-mer value, we suggest experimenting with longer k-mers in the model building phase. All subsequent analyses in this article are performed with a *k*-mer length of 13, unless specified otherwise.

**Fig 2.**
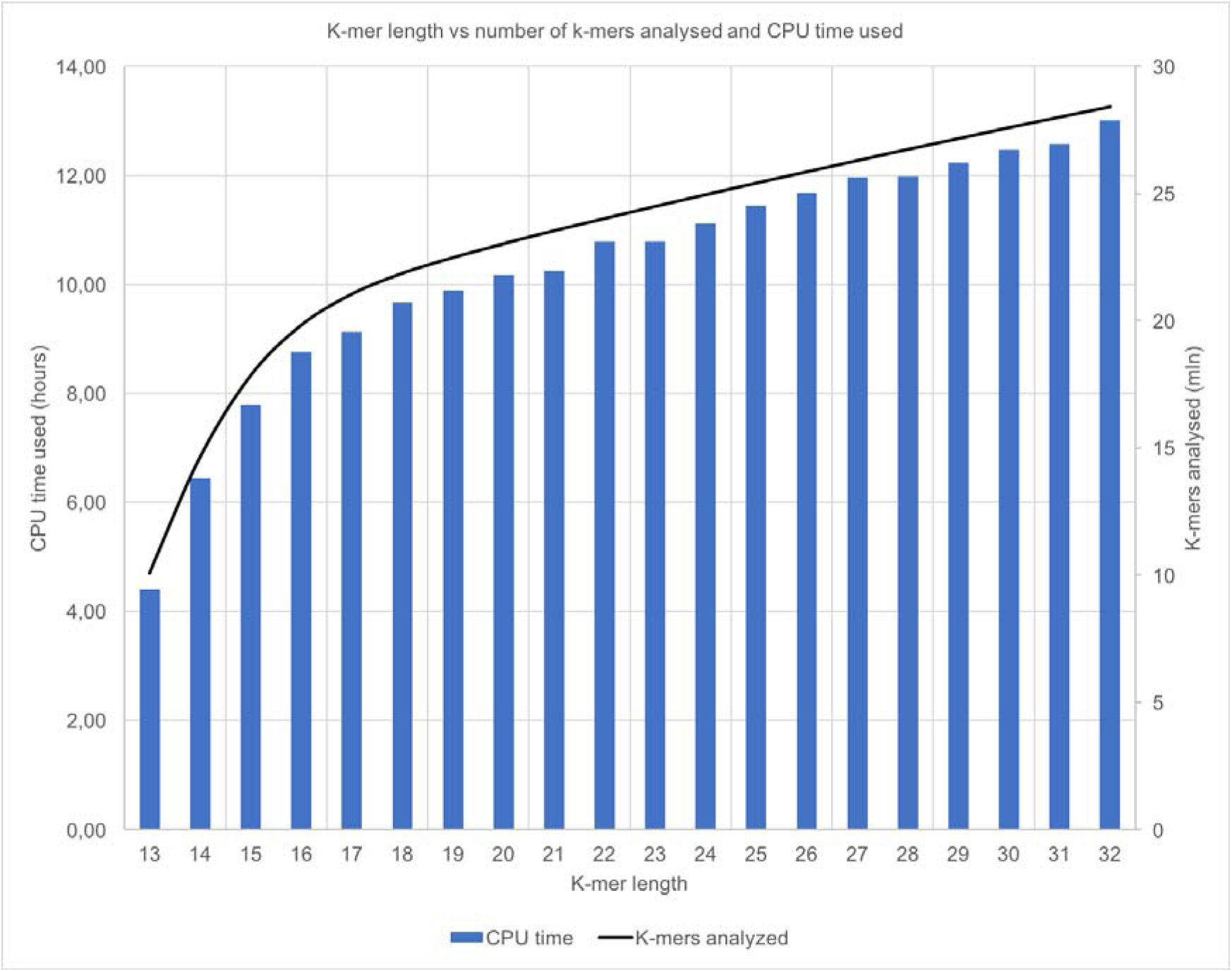
The influence of *k*-mer length on the CPU time of PhenotypeSeeker (bars, left axis) and on the number of different *k*-mers present in the genomes (line, right axis).

### Ciprofloxacin resistance phenotype in *Pseudomonas aeruginosa*

PhenotypeSeeker was applied to the datasets composed of *P. aeruginosa* genomes and corresponding ciprofloxacin MIC-s. We built two separate models using a continuous phenotype for one and binary phenotype for another. Binary phenotype values were created based on EUCAST ciprofloxacin breakpoints [18]. Both models detected *k*-mers associated with mutations in quinolone resistance determining regions (QRDR) of the *parC* (c.260C>T, p.Ser87Leu) and *gyrA* (c.248C>T, p.Thr83Ile) genes (Fig 3, S2 File). These genes encode DNA topoisomerase IV subunit A and DNA gyrase subunit A, the target proteins of ciprofloxacin [19]. Mutations in the QRDR regions of these genes are well-known causes of decreased sensitivity to quinolone antibiotics, such as ciprofloxacin [20]. The model built using a binary phenotype had a prediction accuracy of 88%, sensitivity of 90% and specificity of 87% on the test subset. The coefficient of determination (R^2^) of the test subset for the continuous phenotype was 0.413 (S2 File).

**Fig 3.**
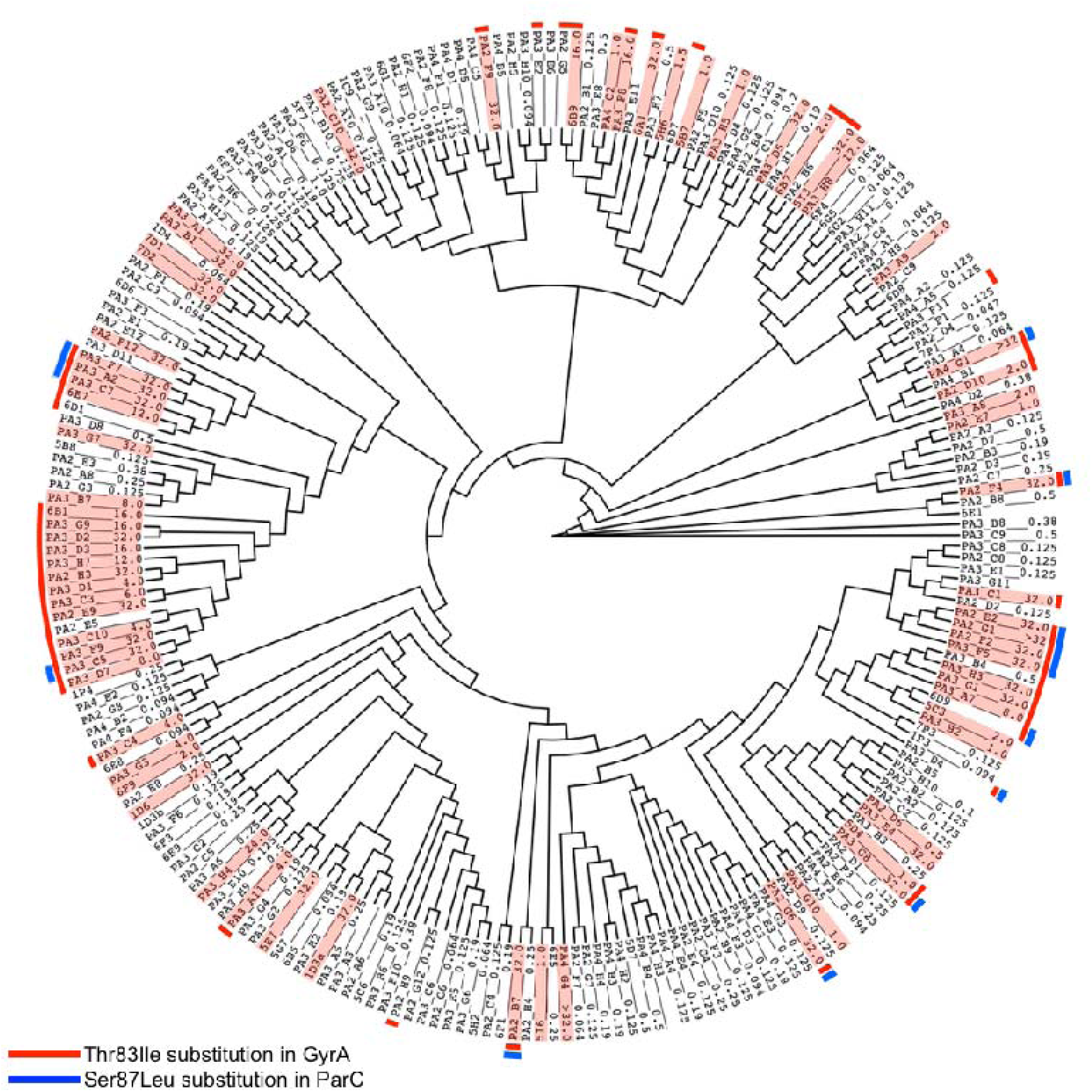
The positions of ciprofloxacin-resistant *P. aeruginosa* strains on cladogram. The MIC values (mg/l) are marked to the external nodes with corresponding strain names. Strains with MIC > 0.5 mg/l are highlighted with pink to denote ciprofloxacin resistance according to EUCAST breakpoints [18]. Strains with detected mutations in QRDR of *gyrA* and *parC* are marked with the color code on the perimeter of the cladogram.

### Azithromycin resistance phenotype in Clostridium difficile

In addition to the *P. aeruginosa* dataset, we tested a *C. difficile* azithromycin resistance dataset (S2 File) studied using Kover in Drouin et al., 2016 [12]. *ermB* and Tn6110 transposon were the sequences known and predicted to be important in an azithromycin resistance model by Kover [12]. *ermB* was not located on the transposon Tn6110. PhenotypeSeeker found *k*-mers for both sequences while using *k*-mers of length 13 or 16. Tn6110 is a transposon that is over 58 kbp long and contains several protein coding sequences, including 23S rRNA methyltransferase, which is associated with macrolide resistance [21]. The predictive models with all tested *k*-mer lengths (13, 16 and 18) contained *k*-mers covering the entire Tn6110 transposon sequence, both in protein coding and non-coding regions. In addition to the 23S rRNA methyltransferase gene, *k*-mers in all three models were mapped to the recombinase family protein, sensor histidine kinase, ABC transporter permease, TlpA family protein disulfide reductase, endonuclease, helicase and conjugal transfer protein coding regions. The model built for the *C. difficile* azithromycin resistance phenotype had a prediction accuracy of 96.5%, sensitivity of 96% and specificity of 97% on the test subset.

### Virulence phenotype in *Klebsiella pneumoniae*

In addition to antibiotic resistance phenotypes in *P. aeruginosa* and *C. difficile*, we used *K. pneumoniae* human infection-causing strains as a different kind of phenotype example. *K. pneumoniae* strains contain several genetic loci that are related to virulence. These loci include aerobactin, yersiniabactin, colibactin, salmochelin and microcin siderophore system gene clusters [22–26], the allantoinase gene cluster [27], *rmpA* and *rmpA2* regulators [28,29], the ferric uptake operon *kfuABC* [30] and the two-component regulator *kvgAS* [31]. The model predicted by PhenotypeSeeker for invasive/infectious phenotypes included 13-mers representing several of these genes. Genes in colibactin (*clbQ* and *clbO*), aerobactin (*iucB* and *iucC*) and yersiniabactin (*irp1, irp2, fyuA, ybtQ, ybtX*, and *ybtP*) clusters showed the most differentiating pattern between carrier and invasive/infectious strains (Fig 4; S2 File). A 13-mer mapping to a gene-coding capsule assembly protein Wzi was also represented in the model. The model built for *K. pneumoniae* invasive/infectious phenotypes had a prediction accuracy of 88%, sensitivity of 91% and specificity of 78% on the test subset.

**Fig 4.**
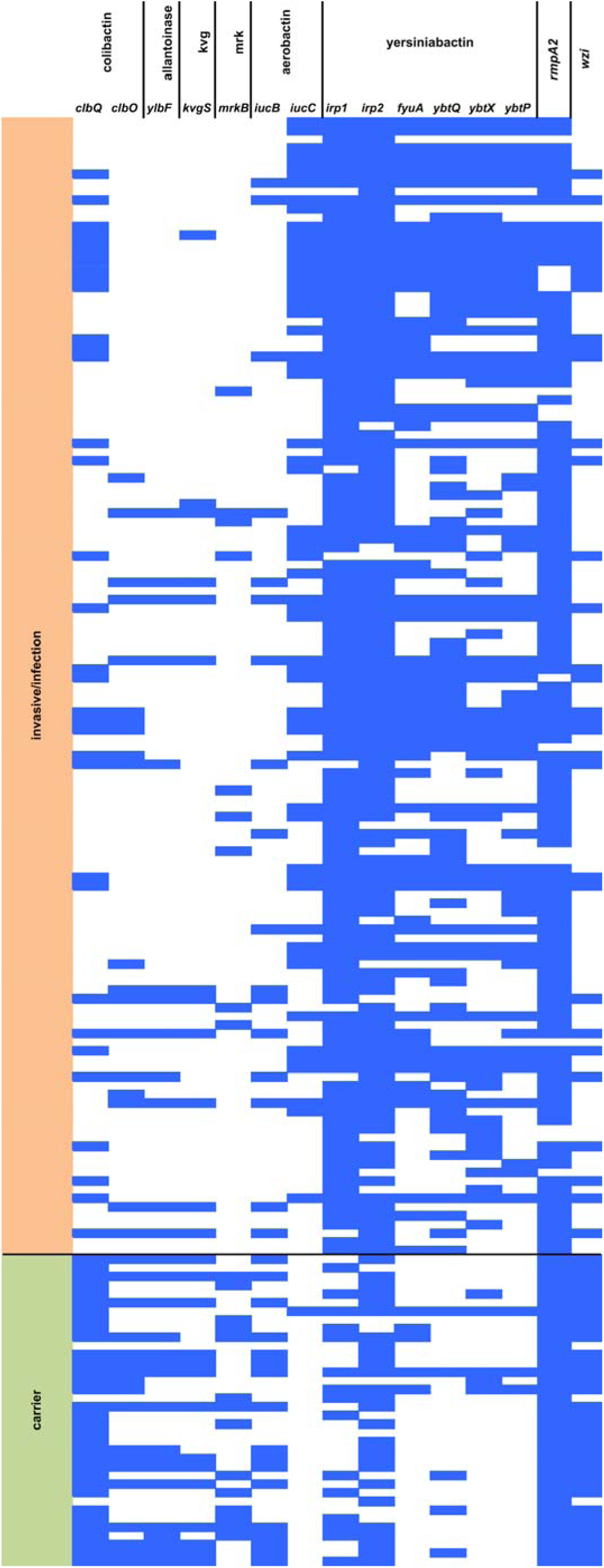
Virulence genes in corresponding clusters and *wzi* included in the PhenotypeSeeker prediction model in *K. pneumoniae* strains (13-mers, weighted, max. 10 000 *k*-mers for the regression model). Each row is one strain, and each column represents one protein coding gene. Blue cells represent 13-mers in the model for the corresponding gene and a strain. Genes in colibactin, aerobactin and yersiniabactin clusters show the most differentiating pattern between carrier and invasive/infectious strains. Virulence genes belonging to the same clusters but without 13-mers in the prediction model are not shown.

### Classification accuracy and running time

To measure the average classification accuracies of logistic regression models, all three datasets were divided into a training and test set of approximately 75% and 25% of strains respectively. A K-mer length of 13 was used, and a weighted approach was tested on binary phenotypes (Table 1). When using sequencing reads instead of assembled contigs as the input, we required a minimum frequency of 5 for a 13-mer to reduce the influence of sequencing errors. The PhenotypeSeeker prediction accuracy is not lower when using raw sequencing reads instead of assembled genomes, and therefore, assembly building is not required before model building. Our results with *K. pneumoniae* show that PhenotypeSeeker can be successfully applied to other kinds of phenotypes in addition to antibiotic resistance.

**Table 1.**
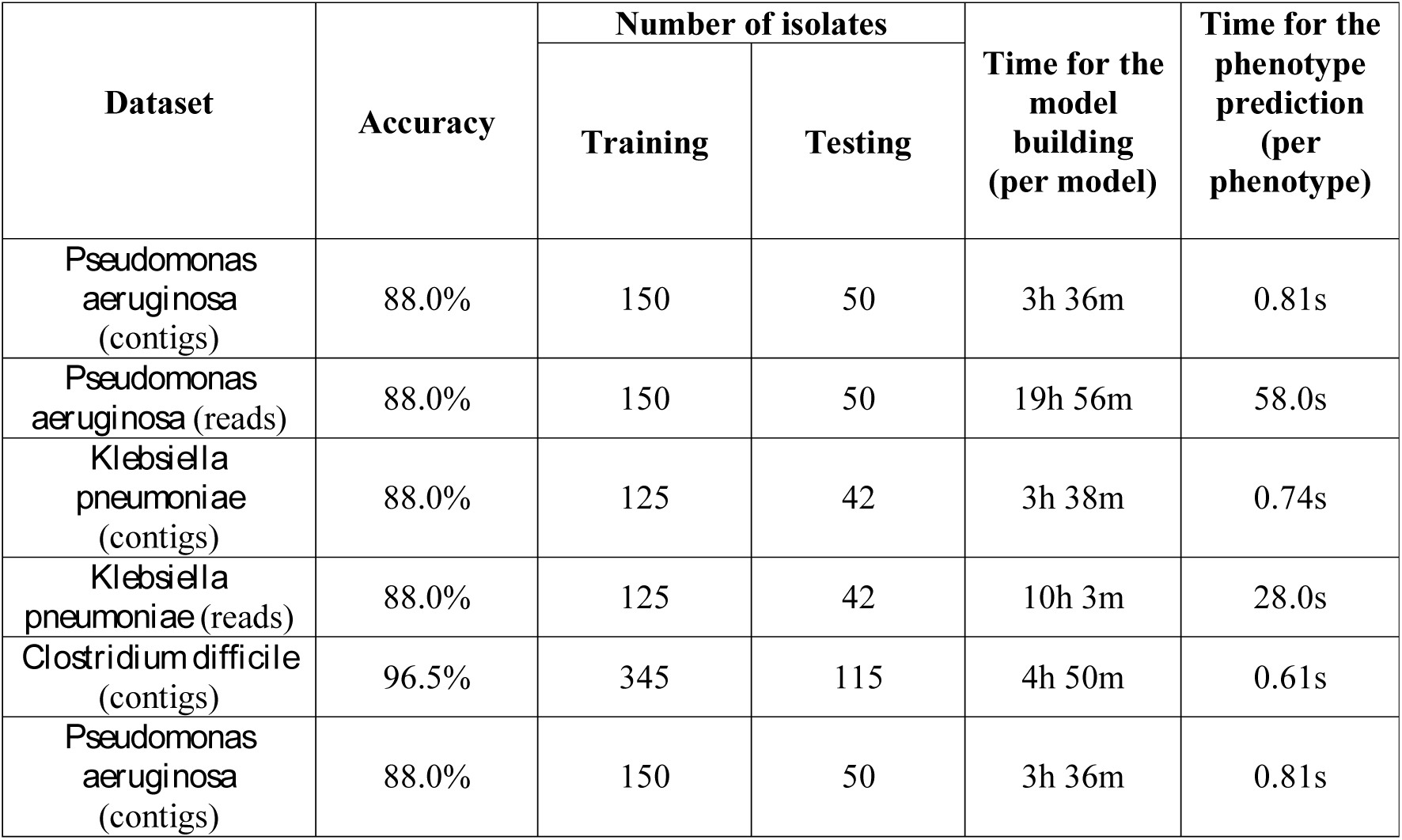
Model prediction accuracy and running time. The results with 13-mers and weighting are shown. The maximum number of 13-mers selected for the regression model was 1000. In cases where sequencing reads were used as the input, a minimum frequency of 5 for a 13-mer was required to reduce the influence of sequencing errors.

In our trials, the model building on a given dataset took 3 to 5 hours per phenotype, and prediction of the phenotype took less than a second on assembled genomes (Table 1). The CPU time of model building by PhenotypeSeeker depends mainly on the number of different *k*-mers in genomes of the training set. The analysis performed on our 200 *P. aeruginosa* genomes showed that the CPU time of the model building grows linearly with the number of genomes given as input (S1 Fig).

The memory requirement of PhenotypeSeeker did not exceed 2 GB if default parameter settings are used, allowing us to run analyses on laptop computers (S2 Fig) if necessary. The p-value cut-offs during the *k*-mer filtering step influence the number of *k*-mers included in the model and have a potentially strong impact on model performance. The tables in the S1 File show the effects of different p-value cut-offs on model performances.

### Comparison with other software

We ran SEER and Kover on the same *P. aeruginosa* ciprofloxacin dataset and *C. difficile* azithromycin resistance dataset to compare the efficiency and CPU time usage with PhenotypeSeeker.

In the *P. aeruginosa* dataset, SEER was able to detect *gyrA* and*parC* mutations only when resistance was defined as a binary phenotype. In cases with a continuous phenotype, those *k*-mers did not pass the p-value filtering step. Since Kover’s aim is to create a resistance predicting model, not an exhaustive list of significant *k*-mers, it was expected that not all the mutations would be described in the output. *gyrA* variation already sufficiently characterized the resistant strains set, and therefore, *parC* mutations were not included in the model. The same applies to the PhenotypeSeeker results with 16- and 18-mers. *parC*-specific 16- or 18-mers were included among the 1000 *k*-mers in the prediction model (based on statistically significant p-values) but with the regression coefficient equal to zero because they were present in the same strains as *gyrA* specific predictive *k*-mers.

In the *C. difficile* dataset, our model included the known resistance gene *ermB* and transposon Tn6110. We were able to find *ermB* with both SEER and Kover. We also detected Tn6110-specific *k*-mers with SEER while running Kover with 16-mers instead of 31-mers as in the default settings.

Regarding the CPU time, PhenotypeSeeker with 13-mers was faster than other tested software programs (3.5 hrs vs 14-15 hrs) without losing the relevant markers in the output (Table 2). Using 16- or 18-mers, the PhenotypeSeeker’s running time increases but is still lower than with SEER and Kover

**Table 2.** PhenotypeSeeker comparison to Kover and SEER using *P. aeruginosa and C. difficile* data. PhenotypeSeeker with the weighting option and maximum 1000 k-mers for the regression model was used.

|  |  | *Pseudomonas aeruginosa* (200 genomes) |  |  | *Clostridium difficile* (460 genomes) |  |  |
| --- | --- | --- | --- | --- | --- | --- | --- |
|  |  | Previously known CIP resistance mutations detected |  |  | Previously known AZM resistance genes* detected |  |  |
| Software | k-mer length | *gyrA* c.248C>T | *parC* c.260C>T | Time for model building | *ermB* | Tn6110 transposon | Time for model building |
| Phenotype Seeker | 13 | + | + | 3h 36m | + | + | 4h 47m |
| Phenotype Seeker | 16 | + | - | 6h 51m | + | + | 9h 7m |
| Phenotype Seeker | 18 | + | - | 7h 31m | - | + | 9h 58m |
| Kover | 16 | + | - | 14h 14m | + | + | 14h 10m |
| Kover | 31 | + | - | 14h 46m | + | - | 13h 40m |
| SEER | 9-100 | + | + | 15h 7m | + | + | 15h 32m |

\* As reported in Drouin *et al.* 2016 [12]

## Discussion

PhenotypeSeeker works as an easy-to-use application to list the candidate biomarkers behind a studied bacterial phenotype and to create a predictive model. Based on *k*-mers, PhenotypeSeeker does not require a reference genome and is therefore also usable for species with very high intraspecific variation where the selection of one genome as a reference can be complicated.

PhenotypeSeeker supports both discrete and continuous phenotypes as inputs. In addition, this model takes into account the population structure to highlight only the possible causal variations and not the mutations arising from the clonal nature of bacterial populations.

Unlike Kover, the PhenotypeSeeker output is not merely a trained model for predicting resistance in a separate set of isolates, but the complete list of statistically significant candidate variations separating antibiotic resistant and susceptible isolates for further biological interpretation is also provided.

Unlike SEER, PhenotypeSeeker is easier to install and can be run with only a single command for building a model and another single command to use it for prediction.

Our tests using PhenotypeSeeker to detect antibiotic resistance markers in *P. aeruginosa* and *C. difficile* showed that it is capable of detecting all previously known mutations in a reasonable amount of time and with a relatively short *k*-mer length. Users can choose the *k*-mer length as well as decide whether to use the population structure correction step. Due to the clonal nature of bacterial populations, this step is highly advised for detecting genuine causal variations instead of strain-level differences. In addition to a trained predictive model, the list of *k*-mers covering possible variations related to the phenotype are produced for further interpretation by the user. The effectiveness of the model can vary because of the nature of different phenotypes in different bacterial species. Simple forms of antibiotic resistance that are unambiguously determined by one or two specific mutations or the insertion of a gene are likely to be successfully detected by our method, and effective predictive models for subsequent phenotype predictions can be created. This is supported by our prediction accuracy over 96% in the *C. difficile* dataset. On the other hand, *P. aeruginosa* antibiotic resistance is one of the most complicated phenotypes among clinically relevant pathogens since it is not often easily described by certain single nucleotide mutations in one gene but rather through a complex system involving several genes and their regulators leading to multi-resistant strains. In cases such as this, the prediction is less accurate (88% in our dataset), but nevertheless, a complete list of *k*-mers covering differentiating markers between resistant and sensitive strains can provide more insight into the actual resistance mechanisms and provide candidates for further experimental testing.

Tests with *K. pneumoniae* virulence phenotypes showed that PhenotypeSeeker is not limited to antibiotic resistance phenotypes but is potentially applicable to other measurable phenotypes as well and is therefore usable in a wider range of studies.

Since PhenotypeSeeker input is not restricted to assembled genomes, one can skip the assembly step and calculate models based on raw read data. In this case, it should be taken into account that sequencing errors may randomly generate phenotype-specific k-mers; thus, we suggest using the built-in option to remove low frequency *k*-mers. The *k*-mer frequency cut-off threshold depends on the sequencing coverage of the genomes and is therefore implemented as user-selectable. One can also build the model based on high-quality assembled genomes and then use the model for corresponding phenotype prediction on raw sequencing data.

## Methods

### Data

PhenotypeSeeker was tested on the following three bacterial species: *Pseudomonas aeruginosa*, *Clostridium difficile* and *Klebsiella pneumoniae.* The *P. aeruginosa* dataset was composed of 200 assembled genomes and the minimal inhibitory concentration measurements (MICs) for ciprofloxacin. The *P. aeruginosa* strains were isolated during the project Transfer routes of antibiotic resistance (ABRESIST) performed as part of the Estonian Health Promotion Research Programme (TerVE) implemented by the Estonian Research Council, the Ministry of Agriculture (now the Ministry of Rural Affairs), and the National Institute for Health Development. Isolated strains originated from humans, animals and the environment (Laht et al., *Pseudomonas aeruginosa* distribution among humans, animals and the environment (submitted); Telling et al., Multidrug resistant *Pseudomonas aeruginosa* in Estonian hospitals (submitted)). Full genomes were sequenced by Illumina HiSeq2500 (Illumina, San Diego, USA) with paired-end, 150 bp reads (Nextera XT libraries) and de novo assembled with the program SPAdes (ver 3.5.0) [32]. MICs were determined by using the epsilometer test (E-test, bioMérieux, Marcy ľEtoile, France) according to the manufacturer instructions. Binary phenotypes were achieved by converting the MIC values into 0 (sensitive) and 1 (resistant) phenotypes according to the European Committee on Antimicrobial Susceptibility Testing (EUCAST) breakpoints [18]. The C. difficile dataset was composed of 460 assembled genomes received from the European Nucleotide Archive [EMBL:PRJEB11776 ((http://www.ebi.ac.uk/ena/data/view/PRJEB11776)] and the binary phenotypes of azithromycin resistance (sensitive=0 vs resistant=1), adapted from Drouin et al., 2016 [11]. The *K. pneumoniae* dataset included 167 isolates analyzed in Holt et al., 2015 [33] using human carriage status vs human infection (including invasive infections) as a binary clinical phenotype (carriage=0 vs invasive/infectious=1). Reads of those 167 strains were de novo assembled with SPAdes (ver 3.10.1) [32]. Therefore, each test dataset was composed of pairs (x, y), where x is the bacterial genome x ∈ {A,T,G,C}^*^, and y denotes phenotype values specific to a given dataset y ∈ {0.008, …, 1024} (continuous phenotype) or y ∈ {0, 1} (binary phenotype).

### Compilation of *k*-mer lists

All operations with *k*-mers are performed using the GenomeTester4 software package containing the glistmaker, glistquery and glistcompare programs [14]. At first, all *k*-mers from all samples are counted with glistmaker, which takes either FASTA or FASTQ files as an input and enables us to set the *k*-mer length up to 32 nucleotides. Subsequently, the *k*-mers are filtered based on their frequency in strains of the training set. By default, the *k*-mers that are present in or missing from less than two samples are filtered out and not used in building the model. The remaining *k*-mers are used in statistical testing for detection of association with the phenotype.

### Weighting

By default, PhenotypeSeeker conducts the clonal population structure correction step by using a sequence weighting approach that reduces the weight of phylogenetically closely related isolates. For weighting, pairwise distances between genomes of the training set are calculated using the free alignment software Mash [15]. Distances estimated by Mash are subsequently used to calculate weights for each genome according to the algorithm proposed by Gerstein, Sonnhammer and Chothia [16]. The calculation of GSC weights is conducted using the PyCogent python package [34]. The GSC weights are taken into account while calculating Welch two-sample t-tests or chi-squared tests to test the *k*-mers’ associations with the phenotype. Additionally, the GSC weights can be used in the final logistic regression or linear regression (if Ridge regularization is used) model generation.

### Chi-squared test

In the case of binary phenotype input, the chi-squared test is applied to every *k*-mer that passes the frequency filtration to determine the *k*-mer association with phenotype. The null hypothesis assumes that there is no association between *k*-mer presence and phenotype. The alternative hypothesis assumes that the *k*-mer is associated with phenotype. The chi-squared test is conducted on these observed and expected values with degrees of freedom=1, using the scipy.stats Python package [35]. If the user selects to use the population structure correction step, then the weighted chi-squared tests are conducted according to the previously published method [36].

### Welch two-sample t-test

In the case of continuous phenotype input, the Welch two-sample t-test is applied to every *k*-mer that passes the frequency filtration to determine if the mean phenotype values of strains having the *k*-mer are different from the mean phenotype values of strains that do not have the *k*-mer. The null hypothesis assumes that the strains with a *k*-mer have different mean phenotype values from the strains without the *k*-mer. The alternative hypothesis assumes that the means of the strains with and without the *k*-mer are the same. The t-test is conducted with these values using the scipy.stats Python package [35], assuming that the samples are independent and have different variance. If the user selects the population structure correction step, then the weighted t-tests are conducted [36]. In that case, the p-value is calculated with the function scipy.stats.t.sf, which takes the absolute value of the t-statistic and the value of degrees of freedom as the input.

### Regression analysis

To perform the regression analysis, first, the x features matrix of the samples is created. The samples in this matrix are strains given as the input and the features represent the *k*-mers that are selected for the regression analysis. The values (0 or 1) in this matrix represent the presence or absence of a specific *k*-mer in the specific strain. The target variables of this regression analysis are the resistance values of the strains. Thereupon, input data are divided into training and test sets whose sizes are by default 75% and 25% of the strains, respectively. In the case of a continuous phenotype, a linear regression model is built, and in the case of a binary phenotype, a logistic regression model is built. In both cases, the Lasso or Ridge regularization can be selected. The Lasso regularization is used by default due to its ability to shrink the coefficients of non-relevant features to zero, which simplifies the identification of k-mers that have the strongest association with the phenotype. To enable the evaluation of the output regression model, PhenotypeSeeker provides model-evaluation metrics. For the logistic regression model quality, PhenotypeSeeker provides the mean accuracy as the percentage of correctly classified instances across both classes (0 and 1). Additionally, the model provides averaged (and for both classes separately) precision, recall and F1-score as a weighted average of precision and recall. For the linear regression model, PhenotypeSeeker provides the mean squared error and the coefficient of determination of the model. To select for the best regularization parameter alpha, a *k*-fold cross-validation on the training data is performed. By default, 25 alpha values spaced evenly on a log scale from 1E-6 to 1E6 are tested with 10-fold cross-validation and the model with the best mean accuracy (logistic regression) or with the best coefficient of determination (linear regression) is saved to the output file. Regression analysis is conducted using the sklearn.linear_model Python package [37].

### Parameters used for training and testing

Our models were created using mainly k-mer length 13 (“-l 13”; default). We counted the *k*-mers that occurred at least once per sample (“-c 1”; default) when the analysis was performed on contigs or at least five times per sample (“-c 5”) when the analysis was performed on raw reads. In the first filtering step, we filtered out the k-mers that were present in or missing from less than two samples (“-min 2 ‐‐max 2”; default) when the analysis was performed on a binary phenotype or fewer than ten samples (“‐‐min 10 ‐‐max N-10”; N – total number of samples) when the analysis was performed on a continuous phenotype. In the next filtering step, we filtered out the *k*-mers with a statistical test p-value larger than 0.05 (“‐‐p_value 0.05”; default).

The regression analysis was performed with a maximum of 1000 lowest p-valued k-mers (“‐‐n_kmers; 1000”; default) when the analysis was done with binary phenotype and with a maximum of 10,000 lowest p-valued *k*-mers (“‐‐n_kmers 10000”; default) when the analysis was performed with a continuous phenotype. For regression analyses, we split our datasets into training (75%) and test (25%) sets (“-s 0.25”; default). The regression analyses were conducted using Lasso regularization (“-r L1”; default), and the best regularization parameter was picked from the 25 regularization parameters spaced evenly on a log scale from 1E-6 to 1E6 (“‐‐n_alphas 25 ‐‐alpha_min 1E-6 ‐‐alpha_max 1E6”; default). The model performances with each regularization parameter were evaluated by cross-validation with 10-folds (“‐‐n_splits 10”; default).

The correction for clonal population structure (“‐‐weights +”; default) and assembly of k-mers used in the regression model (“‐‐assembly +”; default) were conducted in all our analyses.

### Comparison to existing software

SEER was installed and run on a local server with 32 CPU cores and 512 GB RAM, except the final step, which we were not able to finish without segmentation fault. This last SEER step was launched via VirtualBox in ftp://ftp.sanger.ac.uk/pub/pathogens/pathogens-vm/pathogens-vm.latest.ova. Both binary and continuous phenotypes were tested for *P. aeruginosa* and the binary phenotype in *C. difficile* cases. Default settings were used. Kover was installed on a local server and used with the settings suggested by the authors in the program tutorial.

## Acknowledgements

The authors are grateful to Triinu Kõressaar for her invaluable suggestions toward improvement of the manuscript.

## Supporting information

**S1 File. The effects of different p-value cut-offs on model performances.** (PDF)

**S2 File. Phylogenetic trees and isolate specific information of the studied *P. aeruginosa, C. difficile* and *K. pneumoniae* isolates.** (XLSX)

**S1 Fig.**
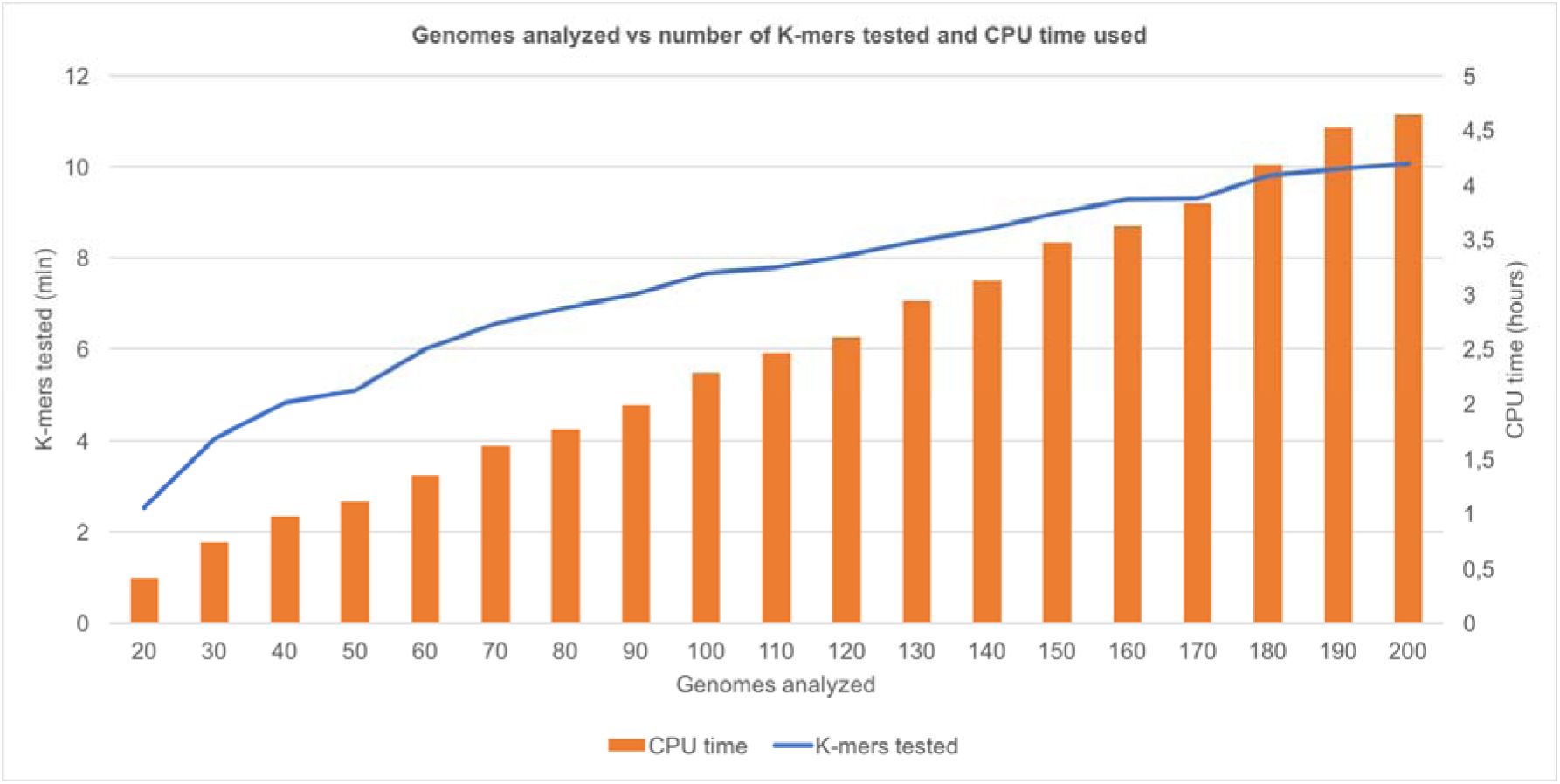
Relationship between the number of input genomes and the CPU time. The PhenotypeSeeker CPU time depends mainly on the number of different k-mers in input genomes and on computations made with every genome. The analysis performed on our 200 P. aeruginosa genomes showed that the PhenotypeSeeker CPU time has a good linear relationship (R2=0.997) with the number of genomes given as input. Although the number of k-mers grows logarithmically with the number of genomes given as input, the linear relationship is because some of the computations made with every genome are more time-consuming when there are larger numbers of different k-mers present in the input genomes.

**S2 Fig.**
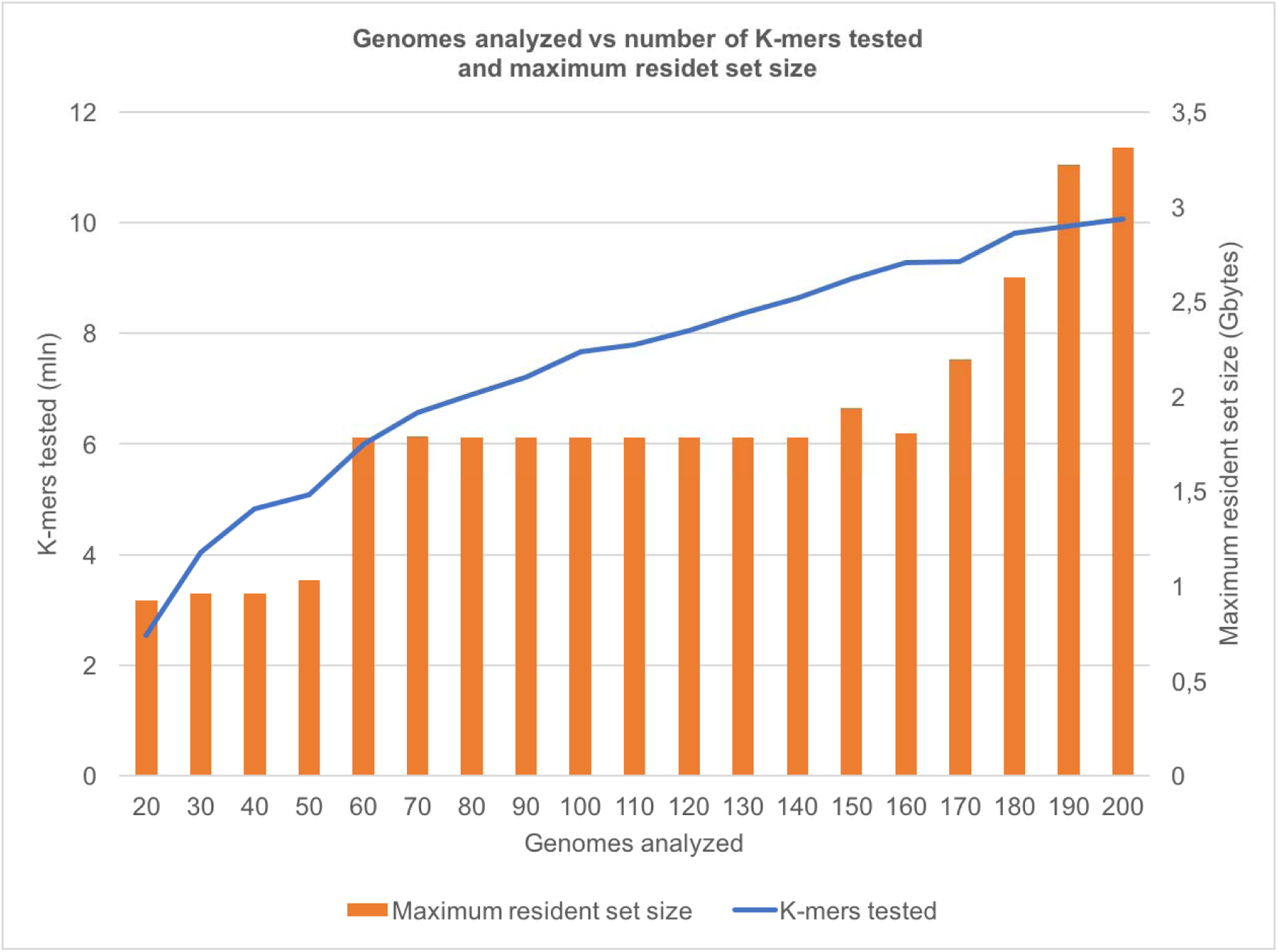
Relationship between the number of input genomes and RAM memory usage. The maximum resident set size of PhenotypeSeeker increases in steps with the number of genomes that are given as the input for model training. This is due to the fact that the maximum resident set size of PhenotypeSeeker is defined by the size of the Python dictionary object into which all different k-mers and their frequencies in genomes are stored. The Python dictionary uses a hash table implementation, and the size of the hash table doubles when it is two thirds full. Therefore, when more genomes are analyzed, more different k-mers are stored into the hash table, and if a certain threshold is exceeded, the next step in the maximum resident set size is taken. However, if the regression is performed with a large number of k-mers, the regression could easily become the most memory using part of the analysis as the data matrix (k-mers x samples), read into memory, grows larger (analysis with 150, 170, 180, 190 and 200 genomes).

